# Adaptation and acclimation of gametophytic traits to heat stress in a widely distributed wild plant along a steep climatic gradient

**DOI:** 10.1101/2024.10.31.621209

**Authors:** Donam Tushabe, Franziska Altmann, Erik Koehler, Sebastian Woods, Sandra Kahl, Sergey Rosbakh

## Abstract

1. Climate change-induced heat waves often result in reduced seed yields and quality via high-temperature effects in the gametophytic phase. Surprisingly, the ability of pollen and ovules, particularly among wild plant populations, to adapt or acclimate to heat stress remains poorly understood.
2. To address this gap, we examined the adaptive and acclimation potential of six gametophytic traits in eleven distinct populations of wild *Silene vulgaris* across a temperature gradient in Europe. First, we cultivated plants in a common garden to reveal differences in gametophytic traits indicative of adaptation. Next, we assessed the acclimation potential of these traits to heat stress by subjecting flowering plants to two chronic heat stress (CHS) treatments: moderate (35/30 °C) and severe (40/35 °C), for 18 days.
3. Findings from the common garden experiment indicated no intraspecific variation in gametophytic traits across the temperature gradient, suggesting that these traits may not influence the plant’s sexual adaptation to its local habitat. Plants originating from colder climates produced more and larger seeds than those from warmer climates. During CHS treatments, the female gametophyte was less temperature sensitive compared to the male gametophyte. Moderate CHS led to larger ovaries with more, large-sized ovules, while severe CHS reduced ovule numbers but increased their size. In contrast, both CHS treatments decreased pollen grain numbers, size, and anther length, with severe CHS causing more significant reductions. These reductions in gametophytic traits ultimately translated to lower seed yield and quality, which may threaten the sustained existence of natural plant populations over time. Under both CHS treatments, the acclimation potential did not vary among plant populations along the temperature gradient, except for pollen size under severe CHS, with larger pollen size in warmer climates than in colder regions.
4. Our findings suggest that the lack of adaptation and acclimation mechanisms in the gametophytic traits (except for pollen size) of wild *Silene vulgaris* populations along the temperature gradient indicates that these plants may rely on alternative strategies, such as shifts in flowering time, to respond to thermal stress.

## Introduction

In the face of rising global temperatures, heat waves have become increasingly severe and frequent, impacting the growth and development of plants (Hatfield and Prueger, 2015). The sensitivity of crucial stages in plant development, such as the gametophytic phase, to heat stress can negatively affect reproductive processes (Hedhly, 2011; Sinha et al., 2021; Yadav et al., 2022). Particularly, the heat-wave effects on pistils, anthers and, therefore, ovules and pollen grains, often leads to diminished seed production and compromised quality (Hatfield and Prueger, 2015; Raza et al., 2019; Tushabe et al., 2023). Consequently, the altered reproductive performance not only jeopardizes food security by causing significant reductions in crop yields (Kumar, 2016; Seppelt et al., 2022) but also carries ecological consequences. As for the former, estimated global crop yields are expected to reduce 10% by 2050 (Wing et al., 2021). As for the latter, altered seed production might cause declines in plant population sizes and disrupt plant-animal interaction across various trophic levels associated with natural populations (Willis et al., 2008; Bogdziewicz et al., 2016).

The ability of species to endure consequences of heat waves mainly relies on either migration to suitable habitats or local adaptation (Pecl et al., 2017; Åkesson et al., 2021). In most cases, migration is not a viable option, due to extensive seed dispersal distances (Ellis, 2015) or lack of suitable new habitats (Rumpf et al., 2018). In this regard, local adaptation and acclimation emerge as crucial survival strategies (Kleine et al., 2021; Wadgymar et al., 2022). While local adaptation involves genetic changes over generations to better suit specific environments (Rehfeldt et al., 2002; Savolainen et al., 2007), acclimation entails short-term physiological adjustments to cope with immediate environmental changes within a plant’s lifetime (Kleine et al., 2021). Both local adaptation and acclimation can enhance heat tolerance through various plant traits. These include maintaining essential physiological processes (e.g., efficient water use or improved photosynthetic mechanisms (Vincent et al., 2020)), phenotypic traits (e.g., early or delayed flowering and or changes in leaf and root morphology (Cook et al., 2012)), and genetic adaptations (e.g., increased production of heat shock proteins (Hasanuzzaman et al., 2013)). As for plant sexual reproduction, previous research has shown that species and populations from climates experiencing frequent and/or severe heat waves tend to have higher heat tolerance of their gametophytic traits (Lancaster and Humphreys, 2020; McDonald et al., 2023; Tushabe and Rosbakh, 2023). Notably, these studies suggest varying tolerance levels, with species from colder regions showing greater cold tolerance, and those from warmer areas displaying higher thermal tolerance (Rosbakh et al., 2016; Lancaster and Humphreys, 2020). Yet, the contribution and relative importance of adaptive changes and acclimation in plant gametophytic responses to heat stress still remains poorly understood. To begin with, existing studies have mainly focused on estimating intra- and interspecific variation of gametophytic traits related to heat tolerance across complex ecological gradients (Di Biase et al., 2021; Kang et al., 2022; Amimi et al., 2023). Although such studies inform us about potential range of plant gametophytic heat tolerance, there is often a challenge in distinguishing the relative contribution of local adaptation and acclimation, because plants usually grow under field conditions that are difficult to control. This limitation can be lifted by cultivating plants representing different populations and species under controlled conditions (i.e., common garden experiment; (Kahl et al., 2019; Tushabe et al., 2023), yet such experimental research has been limited either to cultivated species (see e.g., Hedhly, 2011) or indoor-grown model species (e.g., *Arabidopsis thaliana;* Scheepens et al., 2018). Importantly, just a few of them have considered ovule and pollen traits (e.g., (Flores-Rentería et al., 2018), with most of the studies focusing on flowering phenology (e.g., (Scheepens and Stöcklin, 2013; Arnold et al., 2022) and/or seed traits (e.g., Zhou et al., 2021; Amimi et al., 2023). As a result, our understanding of adaptability and acclimation of gametophytic traits, which are the most sensitive to temperature stress (Hedhly, 2011), in wild plant populations is still lacking.

Here, we close these gaps by testing the adaptive and acclimation potential of six gametophytic traits measured to heat stress in eleven populations of wild *Silene vulgaris* (bladder campion), occurring along a steep climatic gradient of temperature in Europe. In the first part of our experimental study, we cultivated plants in a common garden, to reveal potential long-term adaptations in the gametophytic traits to the local growing conditions. We hypothesized that plants originating from colder climates tend to produce smaller anthers and ovaries with fewer and smaller-sized pollen and ovules, respectively (*H_1_*). The relatively smaller investment into gametophytic tissues of cold-adapted plants should be therefore reflected in lower seed production, both in terms of size and number, compared to their counterparts from warmer climates. This hypothesis is based on the assumption that plant regeneration is constrained in cold environments due to the resource-intensive nature of flowers, gametophytes, and seeds, acting as resource sinks (e.g., (Obeso, 2002).

Next, we investigated the acclimation potential of the same gametophytic traits across the study populations by exposing the flowering plants to experimental chronic heat stress (CHS) treatments: moderate (35/30 °C) and severe (40/35 °C), both lasting 18 days. The moderate CHS treatment represents typical heat waves that plants, especially those in the southern regions, may have already encountered (Table 1; Lhotka and Kyselý, 2022). In contrast, the severe stress represents potential future heat waves that the plants may not have acclimated or adapted to yet (Lin et al., 2022). In response to moderate CHS treatment (35/30 °C), we expected that gametophytic traits in plant populations from warmer climates, experiencing more frequent heat waves, would exhibit a better ability to acclimate (*H_2_*). This is attributed to the presence of acclimation mechanisms developed in response to previous encounters with similar temperatures (e.g., McDonald et al., 2023), enabling these plants to sustain the production of normal-sized anthers, ovaries, pollen, and ovules, resulting in unaltered seed production. Conversely, in plants from colder climates where heat waves are less common, we expected a reduced capacity for acclimation in gametophytic traits due to the lack of previous exposure to such temperatures and the consequent lack of pre-acclimation mechanisms (Nievola et al., 2017). As a result, these plants might produce smaller anthers and ovaries containing fewer and smaller-sized pollen and ovules respectively, potentially resulting in altered seed production and size.

**Table 1:** Origins of the eleven European *Silene vulgaris* populations, including the number of individuals per population (n), the elevation of the g sites, the mean temperature during the flowering period, the mean number of days with temperatures at or exceeding 35 °C and 40 °C, the of flowering period, and the mean annual temperatures in their respective locations. The climatic data are means from 2000 to 2020, derived e European climatic database. The flowering period data are sourced from GBIF occurrence records.

| Population | Country | Site | n | Elevation<br>(m) | Latitude | Longitude | Mean<br>temperature<br>of flowering<br>period (°C) | Mean<br>days<br>≥35<br>(°C) | Mean<br>days<br>≥40<br>(°C) | Flowering<br>period<br>(months) | Mean annual<br>temperature (°C) |
| --- | --- | --- | --- | --- | --- | --- | --- | --- | --- | --- | --- |
| SE80 | Spain | La Palma | 24 | 1100 | 28°46'58.0"N | 17°56'22.0"W | 18 | 0 | 0 | April-June | 14.5 |
| SE83 | France | La Noue du Bourg | 12 | 73 | 46°39'28.0"N | 01°23'13.6"W | 20 | 2 | 0 | April-June | 12 |
| SE84 | Sweden | Vickleby | 12 | 51 | 56°34'37.1"N | 16°27'39.5"E | 20.5 | 0 | 0 | June-August | 7.6 |
| SE49 | Spain | La Gueria Carrocera | 12 | 348 | 43°19'02.0"N | 5°39'45.6"W | 21.3 | 0 | 0 | May-July | 11.4 |
| SE86 | France | Normandie | 12 | 83 | 49°16'06.0"N | 01°37'39.0"E | 22.2 | 3 | 1 | May-July | 10.3 |
| SE91 | Germany | Konstanz | 13 | 400 | 47°40'42.5"N | 9°10'30.0"E | 24 | 4 | 0 | May-July | 9.2 |
| SE35 | Spain | Madrid | 10 | 621 | 40°24'55.0"N | 3°42'45.2"W | 24.5 | 28 | 2 | April-June | 13.8 |
| SE31 | Czech | Brno | 11 | 209 | 49°11'45.4"N | 16°36'31.8"E | 24.7 | 4 | 0 | May-July | 8.4 |
| SE72 | Spain | La Sentiu de Sió | 10 | 316 | 41°48'18.7"N | 0°52'43.6"E | 25 | 22 | 3 | April-June | 14.4 |
| SE98 | Austria | Innsbruck | 10 | 557 | 47°16'14.6"N | 11°24'39.7"E | 26 | 3 | 0 | June-August | 5.7 |
| SE82 | Spain | Alcanó | 23 | 211 | 41°29'29.4"N | 00°26'27.3"E | 26 | 25 | 2 | April-June | 15 |

Under severe CHS treatment (40/35 °C), we hypothesized that all populations would show reduced acclimation potential (*H_3_*). The rarity of exposure to such extreme stress conditions in these plants hinders the development of acclimation mechanisms. Moreover, the physiological constraints imposed by temperatures at 40 °C including the disruption of cellular membranes, protein denaturation, oxidative stress, and impaired gene regulation further contribute to the inability of plants to acclimate to such extreme temperatures (e.g., Araújo et al., 2013; Hasanuzzaman et al., 2013). This is expected to have negative impacts on gametophytic traits, such as reduced pollen and ovule production, ultimately resulting in reduced seed quantity and quality.

## Methods and materials

### Study species and experimental setup

*Silene vulgaris* (Moench) Garcke (Caryophyllaceae), commonly known as bladder campion, is a short-lived herbaceous perennial plant widely distributed across diverse geographical regions, ranging from subarctic to temperate zones (Rabinowitz et al., 1986). It is indigenous to Europe, Asia, and North Africa, with a global presence due to introductions in different regions (USDA NRCS, 2023). Its primary habitats include open grasslands, meadows, woodland edges, as well as disturbed areas such as roadsides, waste places, and metalliferous soils (Marsden-Jones and Turrill, 1957; Friedrich, 1979). The plant exhibits a clump-forming growth habit, with erect or sprawling stems that can reach heights of up to sixty centimeters.

The genus *Silene* has been widely utilized in ecological research due to its favorable attributes, including ease of cultivation and short life cycles, facilitating experimental investigations (Bernasconi et al., 2009; Blavet et al., 2011; Kahl et al., 2019). *Silene vulgaris* particularly offers a distinctive opportunity to investigate how wild plants locally adapt and vary along thermal gradients (e.g., Kahl et al., 2019).

The seeds were collected from eleven populations of *Silene vulgaris* across locations in Austria, Czech Republic, Germany, France, Spain, and Sweden (see Table 1, Figure 1). Seeds were collected in 2015-2019 from at least 10 open-pollinated plants at each site, with at least 1-meter spacing between plants. The exact locations of the sampling sites were recorded using a GARMIN GPS 72H device (accuracy <15 m; Garmin, Switzerland). These populations represent a climatic gradient of *S. vulgaris* in Europe, which may exhibit plant sexual reproduction adaptations to varying temperature conditions. All selected populations exhibit hermaphroditic flowers, containing both male and female reproductive organs within a single flower, and a short seed-to-maturity period of about seven weeks.

**Figure 1:**
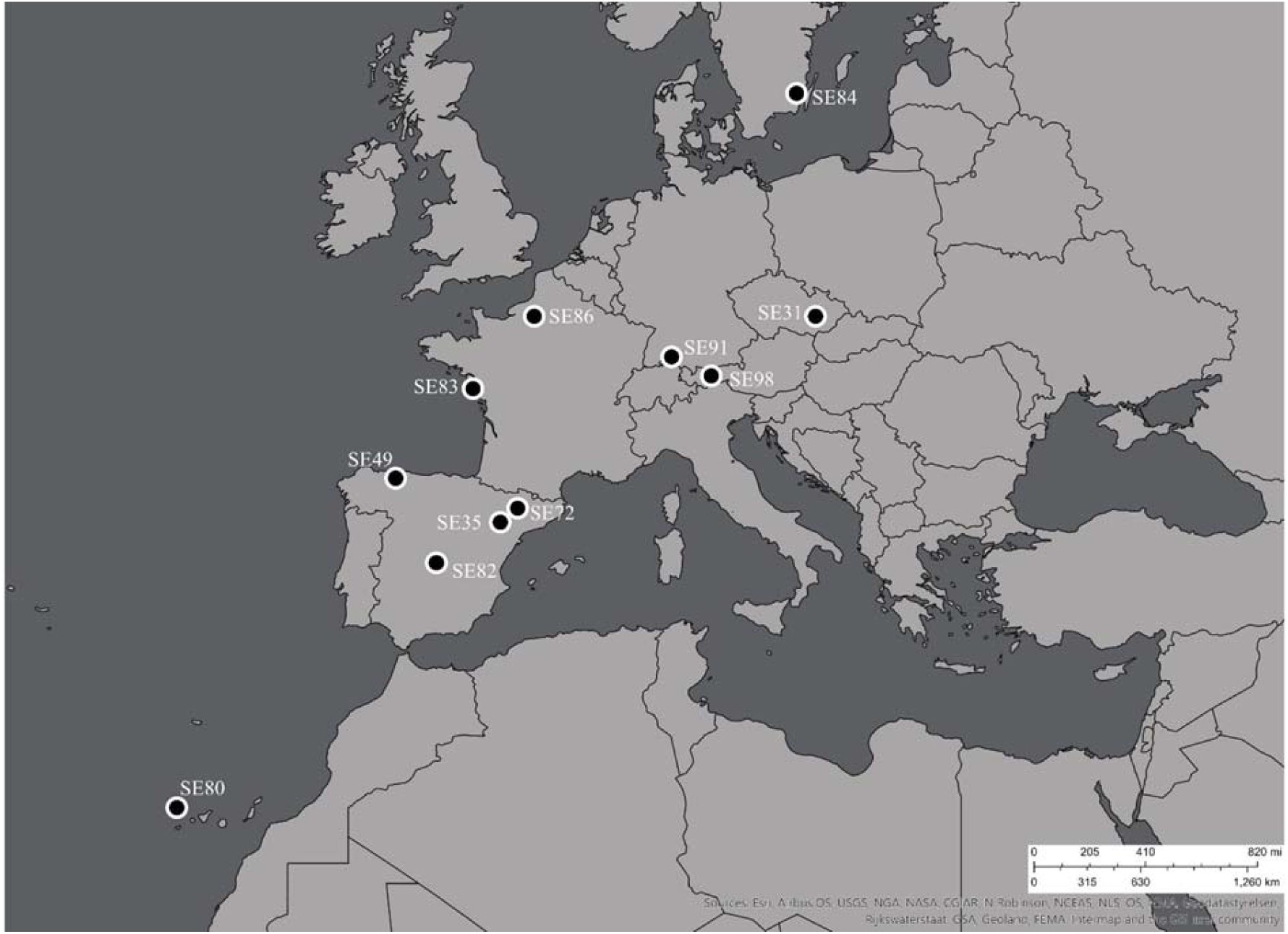
Sampling sites of the eleven *Silene vulgaris* populations across six different European countries. Map created with Esri ArcGIS Online.

The experimental setup and procedures described in this study were adapted from a previous study by Tushabe et al. (2023), conducted within the same greenhouse facility from November 2022 to July 2023.

To eliminate potential maternal effects, we cultivated F1 plants from the seeds collected in the field from each *S. vulgaris* population. These plants were grown in trays filled with a substrate mixture composed of low-nutrient planting soil, coarse sand, and dry compost soil. The plants from distinct populations were bagged before the onset of flowering to prevent cross-pollination.

Thereafter, the F1 seeds were planted in a similar soil substrate as described above. Approximately three weeks after germination, the young plants were individually transplanted into pots, with careful selection based on health and phenological stage to maintain consistency across treatments.

Throughout cultivation, all plants received equal attention, including random rearrangement within the greenhouse and consistent watering. The greenhouse served as a common garden, providing similar conditions (day/night temperatures of 20/15 °C and natural illumination), for investigating the adaptation of gametophytic traits to local growing conditions. Additional lighting was employed to maintain a 12-hour photoperiod during winter months (November to April).

After the emergence of the initial flower bud, plants at similar phenological stage were randomly selected for the CHS experiment.

The CHS experiment was conducted in six identical grow chambers (Homebox Vista Medium, HOMEbox, Germany) equipped with heating mats and LED lamps for temperature and illumination control. Two chambers were maintained at day/night temperatures of 35/30 °C, representing moderate chronic heat stress (CHS), while another two chambers were set at 40/35 °C, indicating severe CHS. Two chambers at 20/15 °C served as the control group. The moderate and severe CHS chambers were used to investigate the acclimation potential of the gametophytic traits, aligning with heat waves currently being observed in European regions and those anticipated in the future (Lin et al., 2022). In each chamber, 10 to 24 plants per population were cultivated for 18 days, ensuring stress treatment at the different developmental stages of pollen and flowers (Mesihovic et al. 2016). To prevent drought stress linked to elevated temperatures, we maintained a consistent soil moisture level by regular watering during the application of heat stress. Post-heat treatment, plants were returned to the greenhouse for seed maturation under controlled conditions.

During cultivation, only the plants in the control group had an aphid infestation and therefore received treatment with Karate zeon (Karate Zeon, Syngenta, Switzerland). Following treatment, the plants were monitored, and no visible changes were observed in the measured traits.

### Measurement of Plant Traits

To evaluate the effects of the experimental treatments on overall plant performance, we conducted measurements of leaf chlorophyll fluorescence, specifically assessing the maximum quantum yield (*Fv/Fm*) of PS II photochemistry. These measurements were conducted over a 20-minute duration using a Pocket PEA tool (Hansatech, Germany) on the final day (day 18) of the treatment period.

### Gametophytic traits

The approach used for gametophytic trait measurements in this study was similar to previous work by Tushabe et al. (2023). Six traits related to male (anther length, pollen production, and size) and female (ovary length, ovule production, and size) gametophytic performance were assessed to investigate the impact of heat treatment on sexual reproduction.

### Seed traits

To evaluate the subsequent impacts of heat stress on gametophytic performance, we measured seed mass and seed production in the treated plants. To accomplish this, the treated plants were kept in the greenhouse for approximately three months until complete seed maturation. During this period, the plants were self-hand-pollinated to ensure the reproductive success of plants. Additionally, the plants were covered with organza bags to prevent any loss of seeds.

Due to low seed production, it was not feasible to collect seeds from individual plants. Instead, all seeds from each accession were combined, and the corresponding seed mass and number were determined. The low seed count may be attributed to limitations in pollination, despite the plants being self-compatible and having undergone hand-pollination.

### Data analysis

Data analyses were conducted using R software version 4.3.0 (R Core Team, 2023).

### Temperature conditions in seed collection sites

To characterize the temperature conditions that each study population experiences during fertilization across the sampled gradient, we calculated the average temperature of the flowering period and identified occurrences of heat waves. To begin with, we extracted species occurrence data per population from the Global Biodiversity Information Facility (GBIF). Our initial assumption was that the GBIF occurrence data were derived from images and herbarium specimens of flowering individuals. Visual inspection of randomly selected images and digitalized herbarium specimens confirmed that assumption at the subsequent stage. Only human observations and preserved specimens with georeferenced locations from the years 2000 to 2020 were considered. This time frame was chosen due to the relatively short lifespan of *Silene vulgaris*, making it more relevant to gather information during the 21st century when significant global climate changes have occurred. The data included details such as the year, month, and days of observation. The flowering period was determined as the month with the highest number of observations, along with the month prior and after it (a total of three months).

Based on the flowering phenology data, we obtained the maximum temperature values for the respective locations within the period from 2000 to 2020. This extraction was performed using the R package *easy climate* (Cruz-Alonso et al., 2023), which relies on high-resolution (1 km) daily climate data sourced from the European climatic database. The mean temperature of the flowering period (MTFP) was computed based on temperatures recorded in the month with the highest number of observations, along with the month prior and after it.

To determine the occurrence of heat waves, we calculated the mean number of days with temperatures at or exceeding 35 °C and 40 °C per accession at the corresponding locations over the period from 2000 to 2020.

### Statistical analysis

To estimate the variability of local adaptation and acclimation potential of the gametophytic traits within study populations across the climatic gradient, we fitted a linear mixed-effects model using the package *lme4* (Bates et al., 2015). The model used one of the measured traits (e.g., chlorophyll fluorescence, gametophyte, or seed trait) as the response variable. The fixed effects included the mean temperature of the flowering period (MTFP), moderate and severe CHS treatments, the control temperature, and their interaction (MTFP*Treatments). To account for individual variability in the response of gametophytic traits, the origin of every plant population accession was included in all models as a random intercept. We visualized how the relationship between mean temperature of the flowering period and gametophytic traits varies across the different heat stress treatments and the control with an interaction plot created using the *interactions* package in R (Long, 2019). Local adaptation was assessed by examining the variability in the control regression slope across the temperature gradient (MTFP). Acclimation was determined by considering both the overall treatment effects and the differences in regression lines between the control and CHS treatments.

Differences in local adaptation and acclimation of gametophytic traits among plant groups from different thermal environments were assessed utilizing the post hoc Tukey test (p<0.05), as applied in the *emmeans* and *multcomp* packages (Hothorn et al., 2008; Lenth, 2023). The chamber effect was not tested in the model because temperature and light conditions were carefully controlled, minimizing potential variations.

## Results

### Leaf chlorophyll fluorescence

The application of both CHS treatments had a significant negative impact on the overall vegetative plant performance regardless plant origin, with the most pronounced effects in severe CHS treatment (mean *Fv*/*Fm* values: control = 0.82, moderate CHS = 0.795, and severe CHS = 0.774; Figure 2, Table 2).

**Figure 2.**
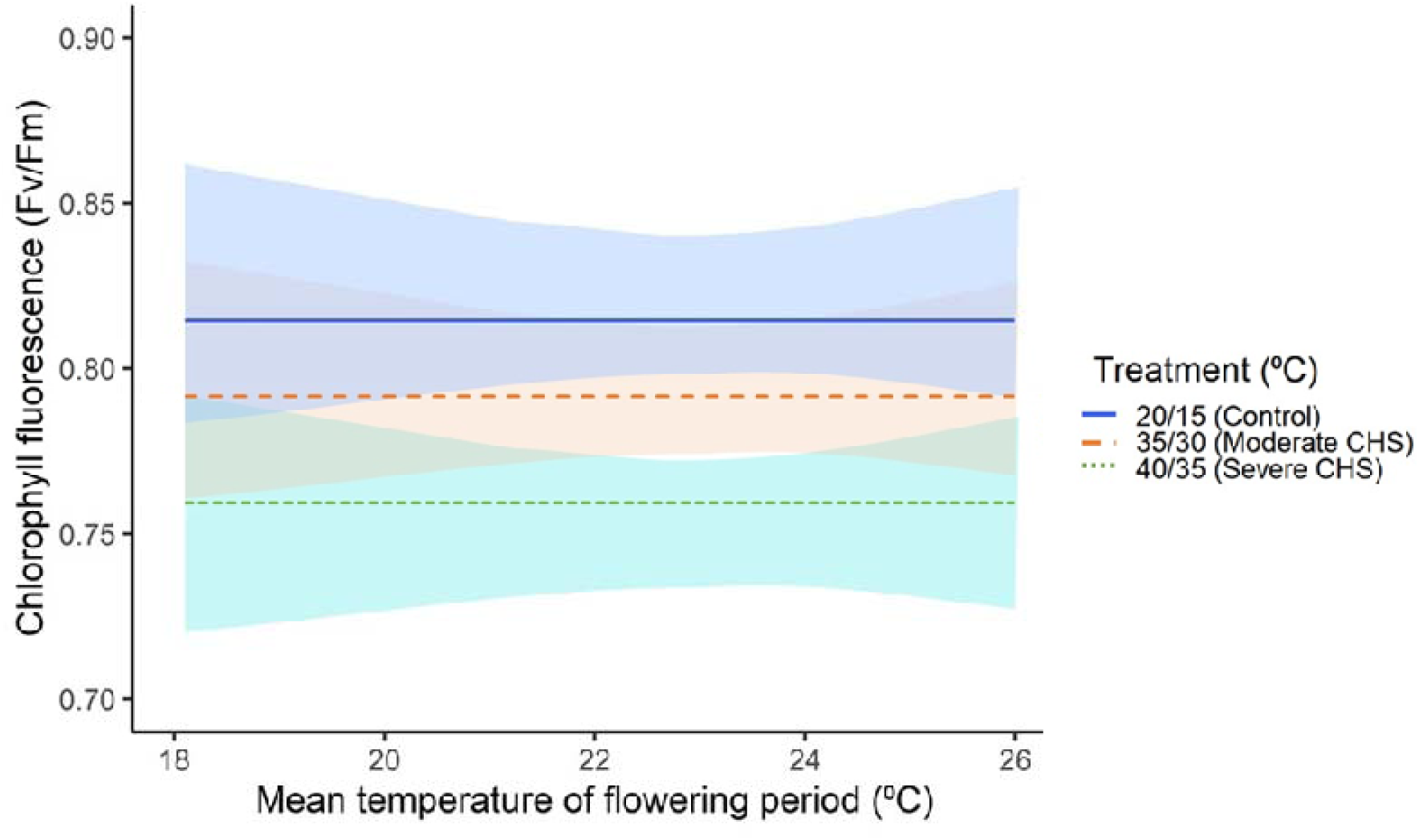
Variation in chlorophyll fluorescence (*Fv/Fm*) values in eleven *Silene vulgaris* populations under two chronic heat stress treatments (moderate and severe), along with a control group, across mean temperature of flowering period. Shaded areas indicate a 95% confidence interval.

**Table 2:** The adaptive (slope of the control regression) and acclimation potential (overall treatment effects and regression slopes) of traits measured (leaf chlorophyll fluorescence [*Fv*/*Fm*], female and male gametophyte, and seed traits) in the eleven wild *Silene vulgaris* populations. Results determined from analyses employing generalized linear mixed-effect models and post-hoc Tukey tests. Significance, denoted by bold formatting, is attributed to treatment/slope effects (p < 0.05). Distinctions between the control and both treatments, as discerned through the Tukey Post-hoc test (p < 0.05), are represented by dissimilar letters. SE refers to standard error.

| Measured traits | Estimates | Control | 35/30°C | 40/35°C | Intercept |
| --- | --- | --- | --- | --- | --- |
| Fv/Fm | Mean | 0.82 a | **0.795 b** | **0.774 c** |  |
|  | SE± | 0.03 | 0.04 | 0.05 | **0.86** |
|  | Slope | -0.002 | 0.0025 | 0.0026 |  |
| Ovary length (mm) | Mean | 3.64 a | **3.91b** | 3.70 ab |  |
|  | SE± | 0.89 | 0.87 | 1.23 | **4.05** |
|  | Slope | -0.023 | -0.037 | -0.061 |  |
| Ovule count | Mean | 88 a | 94 a | **66 b** |  |
|  | SE± | 33.09 | 32.40 | 38.95 | 57 |
|  | Slope | 1.32 | -0.87 | -2.49 |  |
| Ovule size (mm) | Mean | 0.27 a | **0.31 b** | **0.29 bc** |  |
|  | SE± | 0.08 | 0.09 | 0.12 | **0.36** |
|  | Slope | -0.004 | -0.0065 | -0.0029 |  |
| Anther length (mm) | Mean | 2.5 a | **2.2 b** | **2.0 c** |  |
|  | SE± | 0.66 | 0.56 | 0.87 | 2.15 |
|  | Slope | 0.0157 | 0.0433 | -0.0613 |  |
| Pollen count | Mean | 2522 a | **1315 b** | **649 c** |  |
|  | SE± | 878.34 | 832.31 | 583.58 | 2252 |
|  | Slope | 9.35 | -5.62 | -20.13 |  |
| Pollen size (mm) | Mean | 0.043 a | **0.036 b** | **0.033 c** |  |
|  | SE± | 0.01 | 0.01 | 0.01 | **0.04** |
|  | Slope | 4.01E-05 | -3.17E-04 | **0.0015** |  |
| Seed count | Mean | 221 a | **13 b** | **56 c** |  |
|  | SE± | 329.54 | 17.12 | 67.63 | **1894** |
|  | Slope | **-73.55** | -4.88 | -17.51 |  |
| Seed weight (g) | Mean | 0.3 a | **0.02 b** | **0.07 c** |  |
|  | SE± | 0.55 | 0.03 | 0.10 | **3.08** |
|  | Slope | **-0.123** | -0.0067 | -0.026 |  |

### Female gametophytic traits

In the common garden experiment (*H_1_*), there were no differences in female gametophytic traits (ovary length, ovule production, and ovule size) in plants from different climates across the temperature gradient, as the slopes of the corresponding regression lines were not significantly different from zero (Figure 3, Table 2).

**Figure 3.**
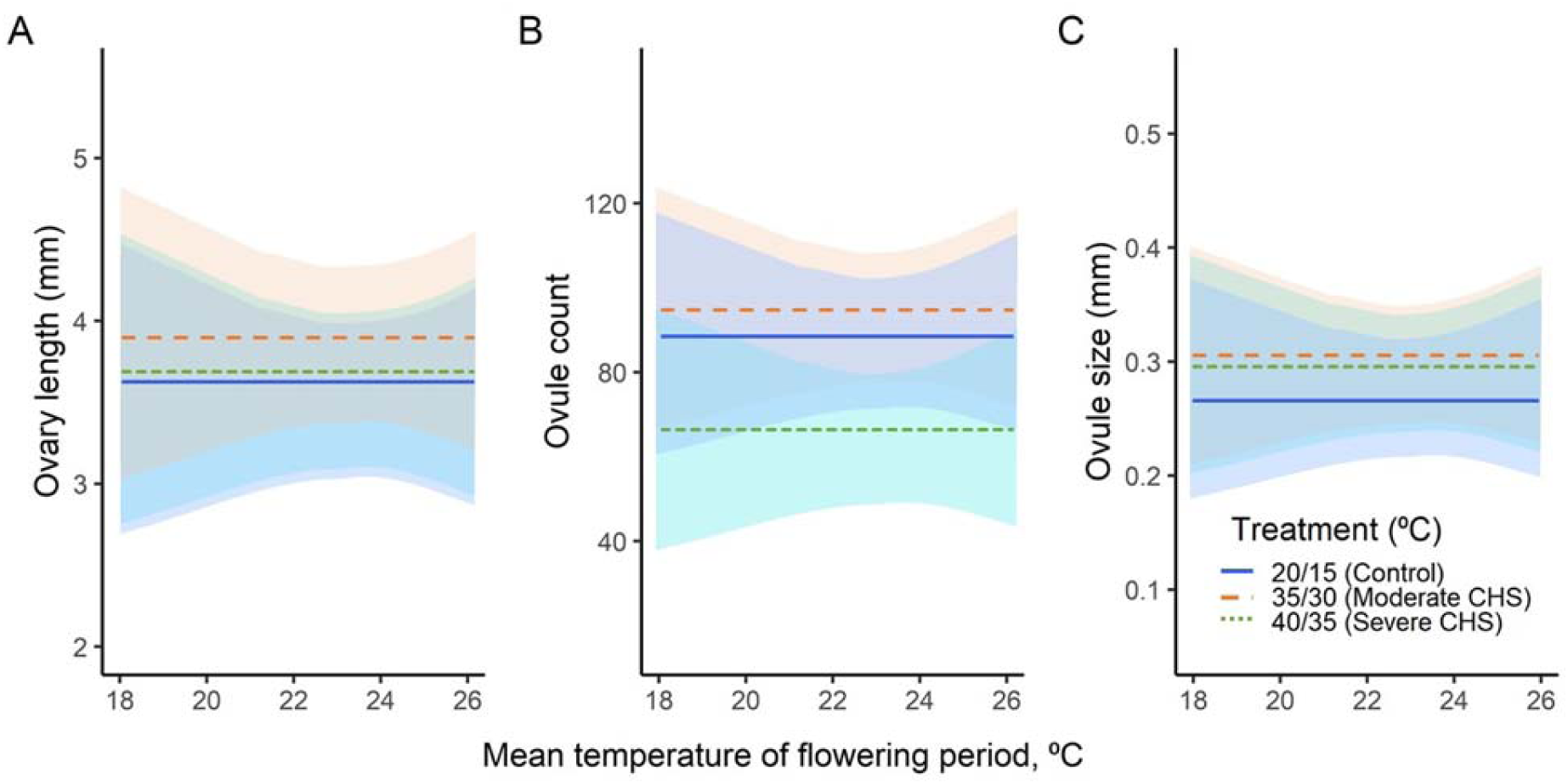
Variations of ovary length (A), ovule production (B), and ovule size (C) in eleven *Silene vulgaris* populations under two chronic heat stress treatments (moderate and severe), along with a control group, across mean temperature of flowering period. Shaded areas indicate a 95% confidence interval. Regression lines parallel to the x-axis indicate lack of statistical differences among the study populations.

When exposed to moderate CHS treatments (*H_2_*), plants produced significantly increased ovary length (3.91 mm) and larger-sized ovules (0.31 mm) compared to the control (3.64 mm and 0.27 mm, respectively; Figure 3A and C; Table 2).

In response to severe CHS treatment (*H_3_*), plants exhibited a significant decrease in ovule production (66 ovules), in contrast to both moderate CHS treatment (94) and the control (88). Contrary, ovule size in the plants significantly increased, as shown by their average ovule size of 0.29 mm compared to the control (0.27 mm; Figure 3B and C; Table 2).

In both moderate and severe CHS treatments, the female gametophytic traits in plants from different climates showed no significant differences along the temperature gradient, as the slopes of the corresponding regression lines were not significantly different from zero (Figure 3, Table 2).

### Male gametophytic traits

In the common garden (*H_1_*), plants originating from both warm and cold climates showed no differences in male gametophytic traits along the temperature gradient, as the slopes of the corresponding regression lines were not significantly different from zero (Figure 4, Table 2).

**Figure 4.**
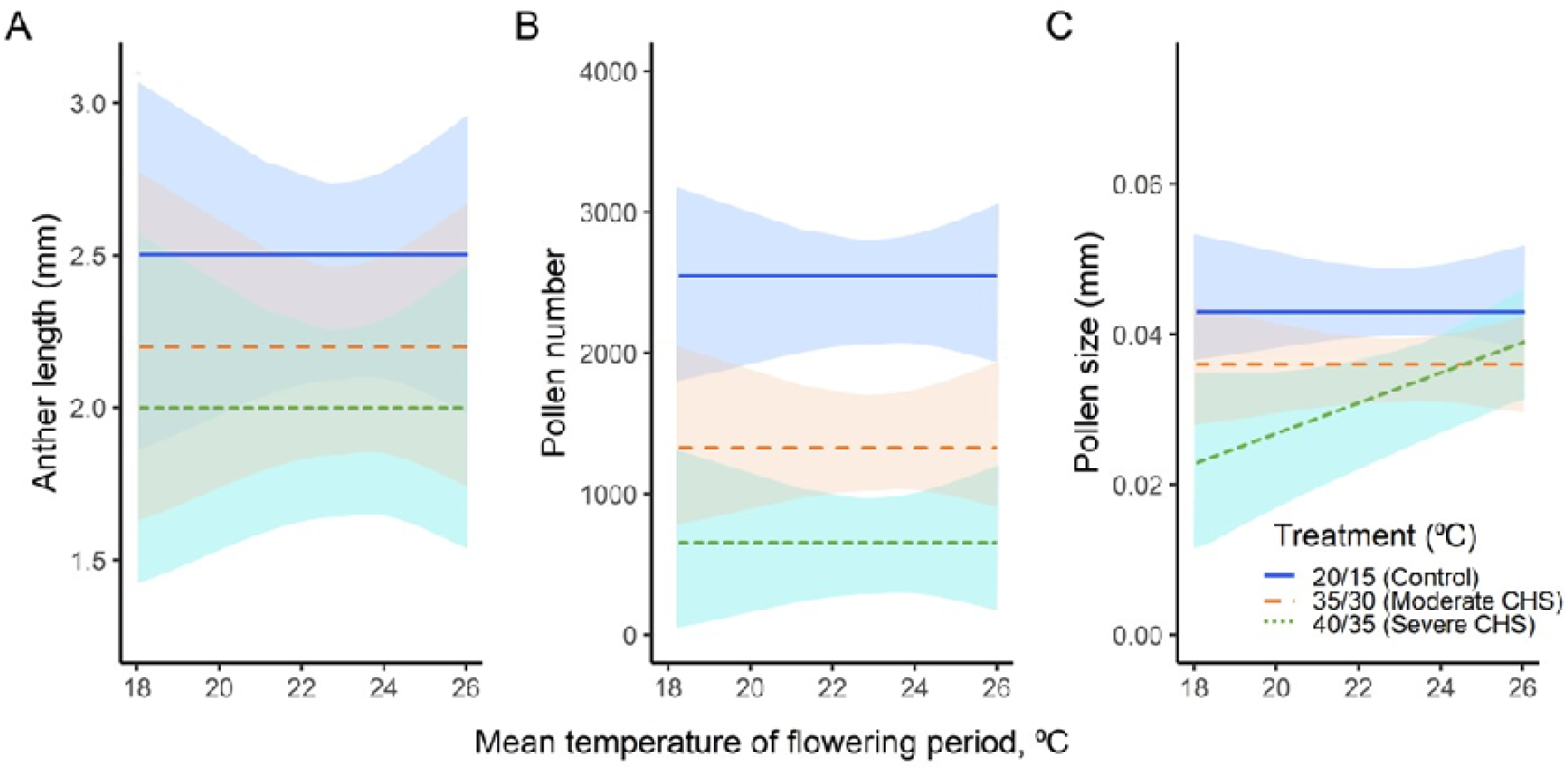
Variations of anther length (A), pollen production (B), and pollen size (C) in eleven *Silene vulgaris* populations under two chronic heat stress treatments (moderate and severe), along with a control group, across mean temperature of flowering period. Shaded areas indicate a 95% confidence interval. Regression lines parallel to the x-axis indicate lack of statistical differences among the study populations.

All three male gametophytic traits measured showed a significant negative response to both heat stress treatments, with significantly larger effect sizes in the severe CHS treatment (*H_3_*) compared to the moderate CHS (*H_2_*) and the control (Figure 4, Table 2). Specifically, in moderate CHS, plants showed significantly reduced anther length (2.2 mm), pollen production (1315), and pollen size (0.036 mm) compared to the control (2.5 mm, 2522 pollen, and 0.043 mm respectively).

Under severe stress conditions (*H_3_*), the male traits further significantly reduced, with the mean anther length decreasing to 2.0 mm, pollen production reducing to 649, and pollen size decreasing to 0.033 mm, in contrast to the control values of 2.5 mm, 2522 pollen, and 0.043, respectively.

Plants originating from different climates did not show any significant differences in both CHS treatments along the temperature gradient, except for pollen size, which exhibited a significant positive correlation with the temperature gradient under severe CHS treatment (p<0.05; Figure 4C, Table 2).

Plants from the warmest climates exhibited larger pollen sizes, while those adapted to colder climates had smaller pollen sizes. Across all plant origins, a 0.0015 mm reduction in pollen size per 1°C decrease in average temperature of flowering period was found under severe CHS.

### Seed traits

In the common garden, plants from warmer climates exhibited a significant negative decline in seed production and mass, with a reduction of 74 seeds and 0.1g (seed mass) per degree of increasing temperature. Conversely, plants originating from colder climates tend to yield more of larger seeds (p<0.05; Figure 5, Table 2).

**Figure 5.**
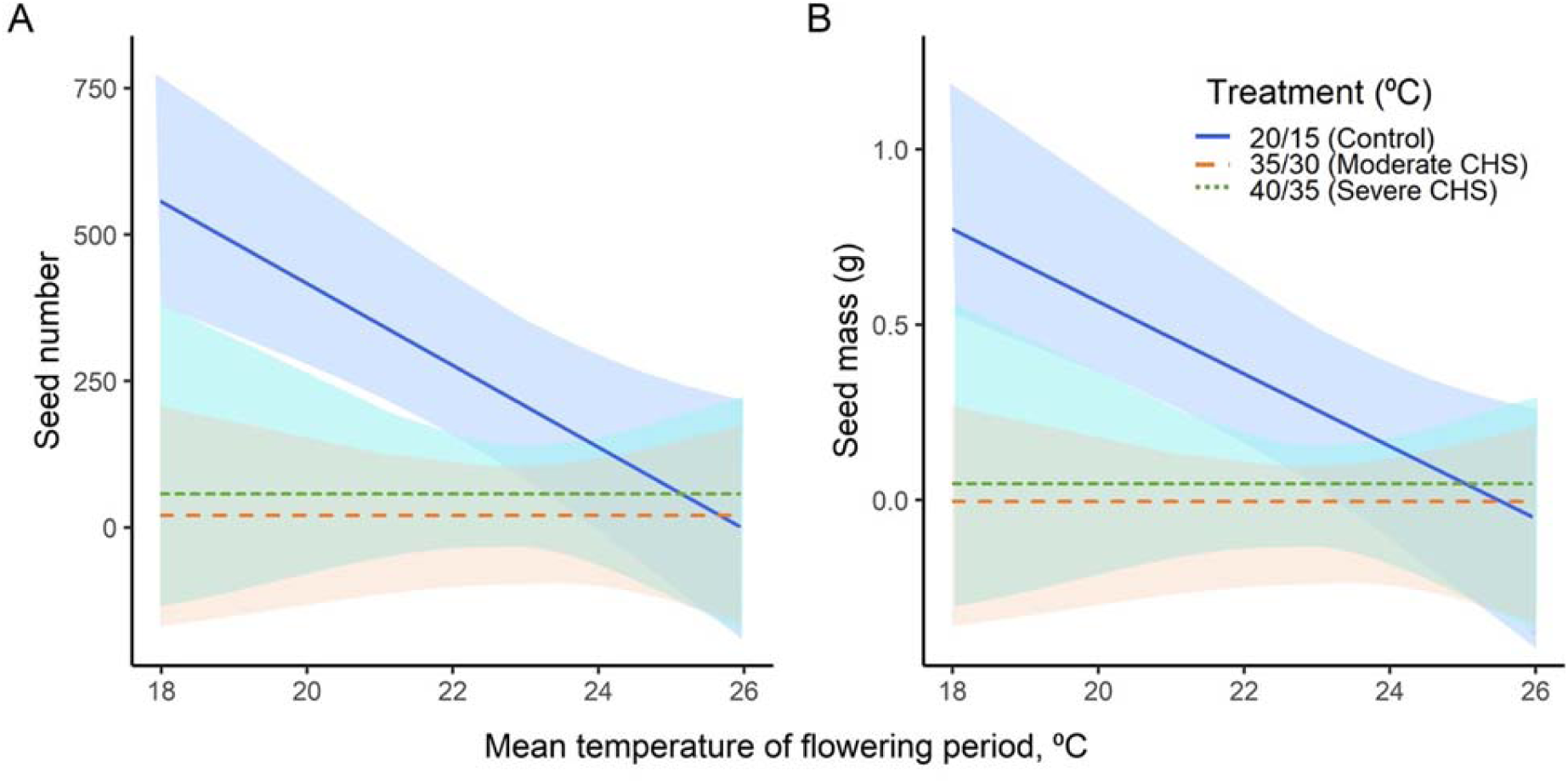
Variations in seed production (A) and seed mass (B) in eleven *Silene vulgaris* populations under two chronic heat stress treatments (moderate and severe), along with a control group, across mean temperature of flowering period. Shaded areas indicate a 95% confidence interval. Regression lines parallel to the x-axis indicate lack of statistical differences among the study populations.

Both CHS treatments applied to the plants resulted in a significant decrease in both mean seed production and seed mass compared to the control group. Specifically, the plants subjected to moderate CHS treatment yielded 13 seeds with a seed mass of 0.02 g, whereas those exposed to severe CHS treatment produced 56 seeds with a seed mass of 0.07 g, in contrast to the control which produced 221 seeds with a seed mass of 0.3 g (Figure 5, Table 2).

Importantly, the resulting seed production and mass from plants across different populations exposed to both CHS treatments showed no differences across the temperature gradient, as the slopes of the corresponding regression lines were not significantly different from zero (Figure 5, Table 2).

Notably, warm-climate populations, characterized by their tendency to yield small seeds in smaller quantities, experienced less pronounced negative impacts on seed production and mass under high temperatures compared to plants in regions with lower temperatures.

## Discussion

### Common garden experiment reveals no intraspecific variation in gametophytic traits (H_1_)

Contrary to our expectations, we found no significant differences in all male and female gametophytic traits measured across the temperature gradient. This finding suggests that these traits in *Silene vulgaris* do not play any role in the plant sexual adaptation to the specific conditions of their local origins. This contradicts the commonly observed variation in other traits such as vegetative traits like canopy height (Jónsdóttir et al., 2023) and specific leaf area (Rosbakh et al., 2015). While research indicates that numerous plant species exhibit local adaptation, some argue that local adaptation might be less prevalent than commonly thought (e.g., Hereford, 2009). For instance, a study by Ebeling et al. (2011) on *Buddleja davidii*, an ornamental shrub, found no evidence of clinal variation in growth and reproductive traits among the different populations.

The lack of clinal variation observed in the gametophytic traits assessed in our study could be attributed to several factors. Firstly, rather than altering the size and number of anthers, ovaries, pollen grains, and ovules, plants may adapt the physiology and biochemistry of these organs. For example, previous studies have shown that warm-adapted pollen tends to have higher concentrations of sugars such as glucose and fructose, more unsaturated fatty acids in lipids, and increased expression of heat-shock proteins and antioxidants to counteract heat stress (Nievola, et al., 2017; Rieu, et al., 2017). On the other hand, cold-adapted pollen tends to have lower sugar levels, increased proportions of saturated fatty acids, and synthesizes cold shock proteins or antifreeze proteins to cope with low temperatures (Satyakam et al., 2022; Jahed et al., 2023). Consequently, these adjustments rarely influence the morphological characteristics of both male and female gametophytes, as could be the case in our study. Secondly, plants possess the ability to adjust their phenological timing to optimize resource utilization under local temperature conditions, thereby minimizing variations in gametophytic traits. For instance, a study on *Arabidopsis lyrata* revealed that southern populations flowered earlier than their northern counterparts (Riihimäki and Savolainen, 2004). Similarly, in our study, populations from southern Europe (e.g., Spain) exhibited early flowering, whereas northern populations (e.g., Sweden) flowered later (see Table 1). This difference could be a strategy for southern populations to evade heat stress associated with high temperatures in later months, while the northern populations may benefit from warmer periods (e.g., Rauschkolb et al., 2023). If plants from diverse locations have adapted to distinct phenological timings in their native environments, it suggests that their reproductive processes are already adjusted to their local conditions.

Lastly, the gametophytic traits might be phylogenetically constrained, i.e., evolutionarily conserved across related species, and thus are not subject to intraspecific variation (e.g., Emilio et al., 2021). Therefore, the lack of clinal variation in our study could be attributed to a combination of physiological/biochemical adaptation, phenological adjustments, and phylogenetic constraints.

Although gametophytic traits did not exhibit clinal variation, significant differences were observed in seed production and mass along the temperature gradient (Figure 5; Table 2). Plants originating from colder climates produced more seeds of greater mass compared to those from warmer climates. This could be attributed to the influence of priority sinks on plant growth patterns, with seeds having the highest sink strength (Wardlaw, 1990; Obeso, 2002). In colder climates, where growth and survival conditions are less favorable, plants tend to allocate more resources towards seed production as an adaptation strategy. This results in the production of larger seeds for better establishment in the unfavorable climatic conditions (Moles and Westoby, 2006; Zhou et al., 2021; Celebias and Bogdziewicz, 2023). On the contrary, within warmer southern populations, plants tend to produce smaller seeds as an adaptation to cope with the stress of higher temperatures (e.g., Zhou et al., 2021). This occurs especially as they approach the edge of their distribution range, where plants are near their tolerance limits. Consequently, these plants allocate fewer resources to seed production, prioritizing other survival mechanisms like efficient water use and growth (e.g., Huot et al., 2014; Lauder et al., 2019; Zhou et al., 2021).

### The acclimation potential of gametophytic traits Overall treatment effects

Acclimation plays a pivotal role in enabling plants to enhance their tolerance to environmental extremes by directly modifying their physiology or morphology (Sumner et al., 2022). The gametophytes of *Silene vulgaris* in our study demonstrated the capacity for acclimation. The results revealed a higher ability to acclimate in female gametophytic traits under both moderate and severe CHS treatments with the male gametophyte exhibiting a reduced capacity to acclimate across both stress treatments.

Female gametophytes exposed to moderate heat stress (*H_2_*), showed larger ovaries, and produced a larger number of larger ovules, whereas male gametophytes produced fewer and smaller pollen grains, with shorter anthers (Figures 3 and 4). The longer ovaries likely provide more space for ovules to develop, and larger ovules may contain more resources, potentially enhancing support for embryo development and increasing the likelihood of viable seed production (e.g., Strelin and Aizen, 2018; Wilkinson et al., 2019). The observed higher ability of female gametophytic traits to acclimate suggests potential trade-offs between female and male traits, where more resources are allocated to female reproductive structures to mitigate the adverse effects of the heat stress on male gametophytes. A study by Gillet and Gregorius (2020) shows that maximizing fertilization success requires more investment in ovule production than in pollen. Moreover, heat stress appeared to directly hinder physiological processes critical for optimal male gametophyte development, leading shorter anthers that produced fewer and smaller pollen grains compared to the control group (Figure 4; Hasanuzzaman et al., 2013; Kumar et al., 2022). Consequently, the reduced number and size of pollen grains decreased the likelihood of successful pollination and fertilization, ultimately resulting in lower seed quantity and quality (Figure 5; Huang et al., 2014; Tushabe et al., 2023).

Averaged over all populations, under severe stress conditions (*H_3_*), fewer but large-sized ovules, less pollen of smaller sizes, with shorter anthers were produced. These alterations in gametophyte morphology likely stem from adjustments in resource allocation to withstand the stress (Ruan et al., 2013; Brock et al., 2017). The fewer yet larger ovules, indicate a shift in resources allocation towards producing fewer but potentially more resilient or viable ovules. This reallocation suggests that larger ovules may have a higher chance of survival or successful fertilization in stressful environments (Gillet and Gregorius, 2020). Conversely, the decrease in pollen production, smaller pollen size, and shorter anther length indicates a reduced investment in male gametophytic structures, likely due to resource scarcity and physiological constraints imposed by severe stress, hindering pollen formation and viability (Müller and Rieu, 2016; Gillet and Gregorius, 2020; Chaturvedi et al., 2021). A similar study showed that male reproduction of *Pinus edulis*, was negatively affected by high temperatures at 40 °C, with stronger effects during pollen germination (Flores-Rentería et al., 2018).

Under both CHS treatments, female gametophytic traits showed greater resilience to short-term heat stress compared to their male counterparts. These finding further confirm the observation that male traits are more susceptible to stress (e.g., Zinn et al., 2010; Chaturvedi et al., 2021; Tushabe et al., 2023). Female gametophytes with thicker ovary tissues are better protected from abiotic stresses (Zinn et al., 2010; Hedhly, 2011), whereas pollen lacks this protection, making it more sensitive to even mild stressors (Bedinger, 1992; Pacini and Dolferus, 2016; Lohani et al., 2020). Furthermore, pollen and tapetum cells show a high demand for energy, indicated by the numerous mitochondria in their cells. Consequently, depletion in energy reserves such as starch and sugar accumulation typically observed under stress conditions, may affect pollen more than other cells (Müller and Rieu, 2016). Additionally, it is proposed that prioritizing the development of the female gametophyte under stressful conditions rather than male gametes could be advantageous (Müller and Rieu, 2016). This strategy could promote outcrossing, leading to greater genetic diversity among progeny and enhancing the chances of genetic adaptation to unfavorable environments (Boyko et al., 2010; Beaudry et al., 2020).

Our study further found that the gametophytic acclimation responses were not effective in coping with both CHS treatments, resulting in a significant decrease in both seed size and number. It is also possible that the heat treatments were too intense for these adaptive mechanisms to counteract the stress. This reduction in ovule and pollen production, resulted in fewer and lower-quality seeds, also reported in previous studies (e.g., Huang et al., 2014; Djanaguiraman et al., 2018). Moreover, the lowered overall performance of the plants, as evidenced by decreased rates of photosynthesis in both heat treatment conditions (Figure 2), likely contributed to the reduced seed production, as the plants’ capacity to generate and store the essential resources vital for seed formation experienced a decline (see, e.g., Sommer et al., 2023). Under high temperature stress, plants may allocate more resources towards growth or survival traits rather than reproduction, potentially enhancing plant fitness (Huot et al., 2014). Supporting evidence for this is found in studies showing a decrease in auxin levels in gametophytes during heat stress, while auxin levels increase in vegetative tissues (Sakata et al., 2010; Sharma et al., 2018).

### Acclimation potential of gametophytic traits along the temperature gradient

All plants from different climates did not show any differences in their acclimation potential to both CHS treatments along the temperature gradient, except for pollen size under severe CHS treatment (Figure 4C). Specifically, plants from warmer climates showed a greater acclimation ability, with larger pollen size in higher temperatures compared to those from colder areas. High temperatures lead to increased evapotranspiration rates, which can rapidly dessicate pollen grains, reducing their viability and thus affecting successful pollination. Therefore, the optimal strategy for plants flowering under high temperatures is to produce fewer but larger pollen grains that are more resilient to volume changes, as observed in our study and supported by Ejsmond et al. (2011). Their study on eight Rosaceae species found that the plants flowering in high temperatures and high evapotranspiration conditions produced significantly larger pollen grains compared to those those in lower temperature and lower evapotranspiration conditions (Ejsmond et al. 2011). Moreover, producing larger pollen could also be a pre-adaptative strategy as larger pollen grains may have increased competitive advantages when interacting with the stigma (Ejsmond et al., 2015). Therefore, warm-adapted plants might prioritize resource allocation towards pollen size rather than pollen number.

The lack of differences in acclimation potential for the other traits across the temperature gradient may be due to similar factors explained in the common garden experiment (see above). These factors include plants prioritizing adaptations in gametophyte physiology and biochemistry over morphological changes in size and quantity (e.g., Nievola, et al., 2017; Rieu, et al., 2017), or adjusting their phenology timings to optimize resource utilization and enhance survival in diverse environments (e.g., Cook et al., 2012; Gugger et al., 2015). Additionally, gametophytic traits might be phylogenetically constrained, thus limiting variations across climatic gradients (e.g., Emilio et al., 2021).

### Conclusion

The evaluation of the male (anther length, pollen production, and size) and female (ovary length, ovule production, and size) gametophytic traits of wild *Silene vulgaris* populations in response to heat stress revealed a lack of adaptation and/or acclimation mechanisms along the temperature gradient. This prompts the question of how gametophytes in natural plant populations, especially in southern regions, cope with heat stress. One plausible explanation is the presence of alternative mechanisms or adaptive strategies, such as alterations in flowering phenology, enabling survival without specific gametophytic adaptations/variations. However, these strategies have limitations due to physiological and, possibly, genetic constraints. For instance, inconsistencies in environmental cues for flowering or sudden environmental changes can disrupt flowering timing, leading to mismatches with optimal gametophytic performance affecting reproductive success (e.g., Fu et al., 2022; Elmendorf and Hollister, 2023). The negative effects on the gametophyte performance can then be translated to reduced seed numbers and quantity, also observed in this study. Producing fewer seeds can lead to reduced dispersal, germination success, and adaptation abilities for plants (Jakobsson and Eriksson, 2000; Soons and Heil, 2002; Long et al., 2015). This could result in smaller, less diverse populations, making species more vulnerable to extinction due to limited abilities to adapt to changing conditions (Long et al., 2015; Schierenbeck, 2017). Consequently, further research into alternative adaptive strategies and mechanisms, including phenotypic plasticity, which was not accounted for in this study, is crucial for gaining insights into the resilience of natural plant populations in the face of ongoing climate change.

## Acknowledgements

We thank the University of Regensburg and the Chair of Ecology and Conservation Biology for the infrastructure and support.

The authors are also grateful to the entire team at the greenhouse of the University of Regensburg for their support throughout the experiment.

## Funding information

Research funding was provided by the Deutsche Forschungsgemeinschaft (DFG), project RO 4909/1-1 (SR). Sergey Rosbakh further appreciates the financial support provided by the NovoNordisk Foundation (Starting grant NNF22OC0078703). Donam Tushabe acknowledges the ‘Bavarian Programme for the Realisation of Equal Opportunities for Women in Research and Teaching’ for the financial support of her PhD programme.

## Declaration of conflict of interest

The authors declare that they have no conflict of interest.

## Data availability statement

The original contributions presented in the study are included in the article/supplementary material; further inquiries can be directed to the corresponding author/s.

## Notes

### Competing Interest Statement

The authors have declared no competing interest.

