## Supplementary Figure S1 for "Adaptation and acclimation of gametophytic traits to heat stress in a widely distributed wild plant along a steep climatic gradient"

### Supplementary material

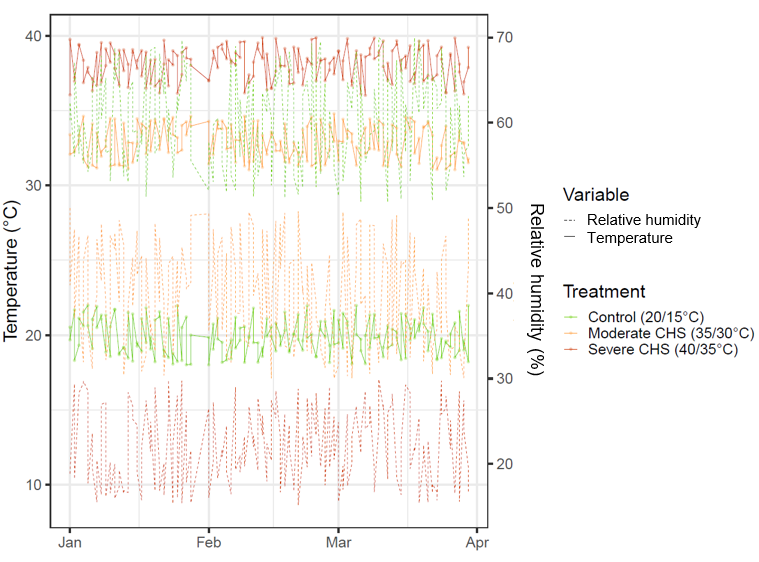

Figure S1: Temperature and relative humidity under control and chronic heat stress treatment conditions during the experiment for the populations of *Silene vulgaris*

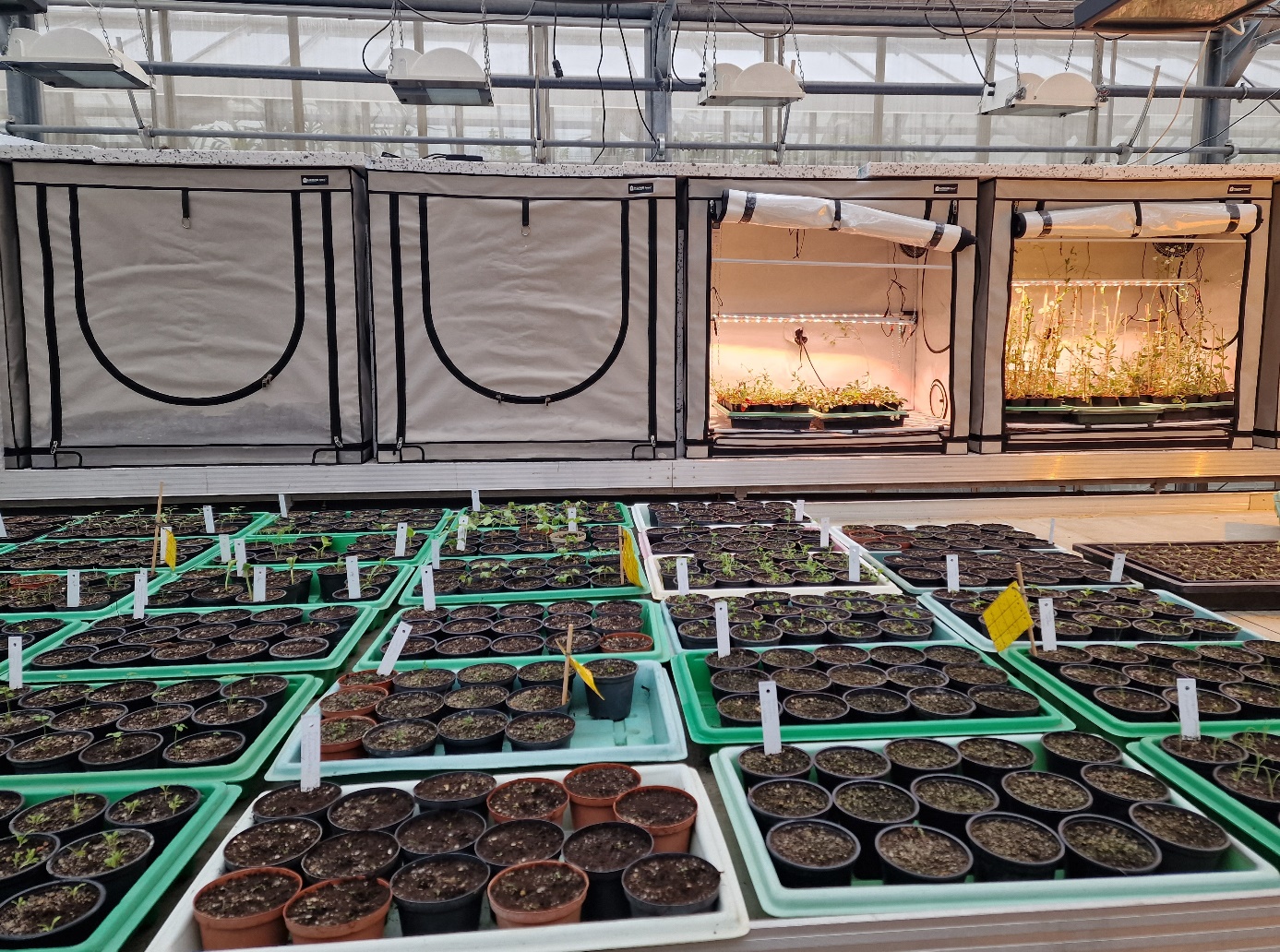

Figure S2: Grow chambers used for the experiment in the greenhouse
